# Genetic architecture underlying HPPD-inhibitor resistance in a Nebraska *Amaranthus tuberculatus* population

**DOI:** 10.1101/2021.06.11.448079

**Authors:** Brent P Murphy, Roland Beffa, Patrick J Tranel

**Author notes:** Correspondence to: PJ Tranel, Department of Crop Sciences, 1201 W. Gregory Dr., Urbana, IL 61801 USA.

## Abstract

**BACKGROUND:** *Amaranthus tuberculatus* is a primary driver weed species throughout the American Midwest. Inhibitors of 4-hydroxyphenylpyruvate dioxygenase (HPPD) are an important chemistry for weed management in numerous cropping systems. Here, we characterize the genetic architecture underlying the HPPD-inhibitor resistance trait in an *A. tuberculatus* population (NEB).

**RESULTS:** Dose-response studies of an F_1_ generation identified HPPD-inhibitor resistance as a dominant trait with a resistance/sensitive ratio of 15.0-21.1. Segregation analysis in a pseudo-F_2_ generation determined the trait is moderately heritable (H^2^ = 0.556), and complex. Bulk segregant analysis and validation with molecular markers identified two quantitative trait loci (QTL), one on each of Scaffold 4 and 12.

**CONCLUSIONS:** Resistance to HPPD-inhibitors is a complex, largely dominant trait within the NEB population. Two large-effect QTL were identified controlling HPPD-inhibitor resistance in *A. tuberculatus*. This is the first QTL mapping study to characterize herbicide resistance in a weedy species.

## 1 INTRODUCTION

*Amaranthus tuberculatus* (Moq.) Sauer (waterhemp) is a summer annual forb native to the American Midwest^1^. As a primary driver weed species within annual production agriculture systems, the control of *A. tuberculatus* is a core component of weed management systems in afflicted regions^2^. If uncontrolled, *A. tuberculatus* can cause severe yield losses (up to 74%) in *Zea mays* L. (maize)^3^. A major contributing factor to the pervasiveness of *A. tuberculatus* as a problematic weed is its ability to evolve herbicide resistance. Resistance to herbicides from seven unique sites of action have been reported in *A. tuberculatus*, with numerous examples of multiple-resistant populations^2,4,5^.

The enzyme 4-hydroxyphenylpyruvate dioxygenase (HPPD; EC 1.13.11.27) catalyzes the second step of a pathway that yields tyrosine, plastoquinone, and tocopherol^6^. This enzyme is the target of herbicides, such as tembotrione, belonging to group 27, and its inhibition leads to photooxidative destruction of chlorophyll and photosynthetic membranes^7^. Resistance to HPPD-inhibitors was first reported in 2009 in *A. tuberculatus* and has since been observed in two other species (*Amaranthus palmeri* S. Wats and *Raphanus raphanistrum* L.)^8^ (Heap I, http://weedscience.org). While the underlying genetic elements responsible for HPPD-inhibitor resistance in weeds has yet to be established, several studies have reported the involvement of cytochrome P450 monooxygenases^8–15^. The genetic architecture underlying the trait is poorly understood beyond simple F_2_ inheritance studies^13,14^ and RNA sequencing studies^15^. In each investigated population, the authors concluded HPPD-inhibitor resistance is due to more than one gene.

Genetic mapping approaches are powerful tools to uncover the genetic architecture of traits of interest. Alleles linked to a trait will co-segregate, allowing the identification of genomic regions that affect the trait^16^. Bulk segregant analysis is an application of traditional quantitative trait loci (QTL) mapping, in which individuals of a segregating population are grouped based on phenotype^17,18^. Recently, genomes for *A. tuberculatus* have been released, enabling high-throughput mapping strategies in this non-model species^19,20^.

In 2010, a 2,4-D resistant *A. tuberculatus* population, termed FS, was identified in southeastern Nebraska^21^. While an initial screening did not identify HPPD-inhibitor resistance^22^, this was conducted at a single rate. Further analysis identified consistent HPPD-inhibitor resistance^23^. Interestingly, the field from where the population was collected has no field-use history of HPPD-inhibitors. The objective of this study was to characterize the genetic architecture underlying HPPD-inhibitor resistance in this population through inheritance, segregation, and linkage mapping.

## 2 MATERIALS AND METHODS

### 2.1 Plant populations and growth conditions

The NEB population (originally FS, renamed on transfer to Illinois) was originally derived from a little bluestem grass (*Schizachyrium scoparium* (Michx.) Nash ‘Camper’) field from Cass County, Nebraska^21^. The sensitive control population WUS originated from the north bank of the Ohio River, Ohio, USA^24^. Seeds were surface sterilized in 50% bleach for 20 minutes, rinsed three times with deionized water, and suspended in 0.1% agarose. Seeds were stratified for one week at 4°C and germinated in sterile Petri dishes on saturated filter paper (Whatman) under greenhouse conditions. Experiments conducted in the greenhouse followed a 12:12 h day:night cycle, with temperatures ranging from 28-30°C during the day and 25-27°C during the night. Experiments conducted in the growth chamber followed a 16:8 day:night cycle, with temperatures ranging from 28-30°C during the day and 25-27°C during the night.

### 2.2 Parent selection

Plants were grown to the 12-leaf stage in the greenhouse, whereupon apical meristems were removed to promote branching. Clones were propagated from branches^23^ and grown to approximately 15 cm in height in the growth chamber to dissuade flowering. Two clones from each of 16 and 12 plants from the NEB and WUS populations, respectively, were screened for their responses to the herbicide tembotrione (Laudis, Bayer Crop Science). Clones of WUS plants were screened for herbicide sensitivity at 0.92 g ai ha^-1^, while clones of NEB plants were screened for herbicide resistance at 14 g ai ha^-1^. Herbicide treatments were applied using an 80015 even flat-fan nozzle (TeeJet Technologies) in a spray chamber, approximately 46 cm above the plant canopy. All tembotrione treatments included 1% v/v crop oil concentrate and 1.875% v/v urea ammonium nitrate. Response to herbicide was visually evaluated three weeks after application, and the most resistant (R) NEB and most sensitive (S) WUS plants were selected. Using plants from which the clones were derived, the following single-plant crosses were conducted in the greenhouse in isolation: RxR, R(female)xS(male), and SxS.

### 2.3 Inheritance and segregation analysis

Dose-response analysis was conducted to characterize trait inheritance. An eight-step, four-fold rate titration was conducted with tembotrione, with treatments ranging from 0.18 to 2941 g ai ha^-1^ and applied to plants at the 8-10 leaf stage in 164 mL cone-tainers. Six plants per population per dose were assayed in each of two experiments separated by time. Above-ground tissue was collected and dried 21 days after application. Biomass was fitted to a four-parameter log-logistic model using the drc package in R^25^. Twenty-four plants of the RxS progeny were intermated to generate a segregating pseudo-F_2_ population. Segregation in the pseudo-F_2_ population was measured in response to tembotrione at 11.5 g ai ha^-1^ across two experimental runs, with 288 plants in run A and 144 plants in run B. At 21 days after application, plants were visually rated as resistant or sensitive and above-ground tissue was collected and dried. Variance from environment and error was calculated as the average variance of SxS and RxR controls grown with the pseudo-F_2_ plants, and broad-sense heritability (H^2^) calculated from the following equation:

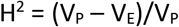

where V_P_ is variance observed among pseudo-F_2_ plants and V_E_ is the variance from environment and error.

### 2.4 Library prep and sequencing

DNA extractions from single leaves of parent and pseudo-F_2_ plants were conducted following the hexadecyltrimethylammonium bromide (CTAB) method^26^. Parent DNA samples were assessed for quality and quantity using a spectrophotometer (Nano Drop 1000 spectrophotometer, Thermo Fisher Scientific), and the whole genomes were shotgun-sequenced using a HiSeq4000, with 150 bp paired-end reads (Illumina). Whole genome sequencing yielded approximately 68x depth for NEB and 65x for WUS. Double-digest restriction site associated DNA sequencing (ddRADseq) libraries were generated from pseudo-F_2_ plants following the protocol optimized for *A. tuberculatus* used by Montgomery et al^27^. Libraries were sequenced using a NovaSeq6000, with 100 bp single-end reads (Illumina). All sequencing was conducted at the Roy J Carver Biotechnology Center at the University of Illinois.

### 2.5 QTL discovery and validation

Whole-genome sequencing reads were aligned to the female reference genome for *Amaranthus tuberculatus^19^* with BWA and variants called using a GATK 4.0 best practices pipeline^28–29^. For each sample, over 98% of reads mapped to the reference genome. The following variant hard filters were established for INDEL (FS > 30, −4 < ReadPosRankSum < 4, SOR > 4, QD < 10) and SNP (FS > 20, −3 < ReadPosRankSum < 4, SOR > 4, QD < 10, MQ < 50), yielding almost 1.6 M mapping-quality variants (homozygous in NEB, absent from WUS). ddRADseq libraries were de-multiplexed and barcodes removed and variants called using a GATK 4.0 best practices pipeline^29^, in which samples were computationally binned by visual rating (sensitive: 60 plants; resistant: 141 plants) prior to the HaplotypeCaller stage. An average of 2.9 M reads per sample were obtained and 148 samples had over 2.5 M reads; approximately 15x coverage based on prior RAD-seq studies in *A. tuberculatus^27^* (data not shown). Bulk segregant analysis was conducted with the QTLseqr package in R^30^, using the QTL-seq method developed by Takagi et al^31^. Putative QTL were considered a stretch of >2 continuous significant windows (α = 0.05). Molecular markers were developed within the intervals of putative QTL based on parental sequencing data and confirmed on parental DNA samples. PCR was conducted using 1x Green GoTaq Flexi Buffer (Promega), 2.5 mM MgCl_2_, 1mM of each dNTP, 0.4 μM of both F and R primers (Table 1), and 0.025 U Bullseye Taq DNA Polymerase (MidSci). When indicated, restriction digests were conducted using the Cutsmart System (New England Biolabs), in which digests occurred overnight at 37°C. PCR was conducted with the following conditions: initial denaturation at 94°C for 30 sec, 40 cycles of 94°C for 10 sec, variable for 15 sec (Table 1), 72°C for 60 sec, and a final extension step at 72°C for 60 sec. PCR and restriction digest products were visualized on 2% agarose gels using gel electrophoresis. A total of 223 pseudo-F_2_ samples were binned by genotype and separation of groups was assayed using Tukey’s HSD from the agricolae package in R.

**Table 1.**
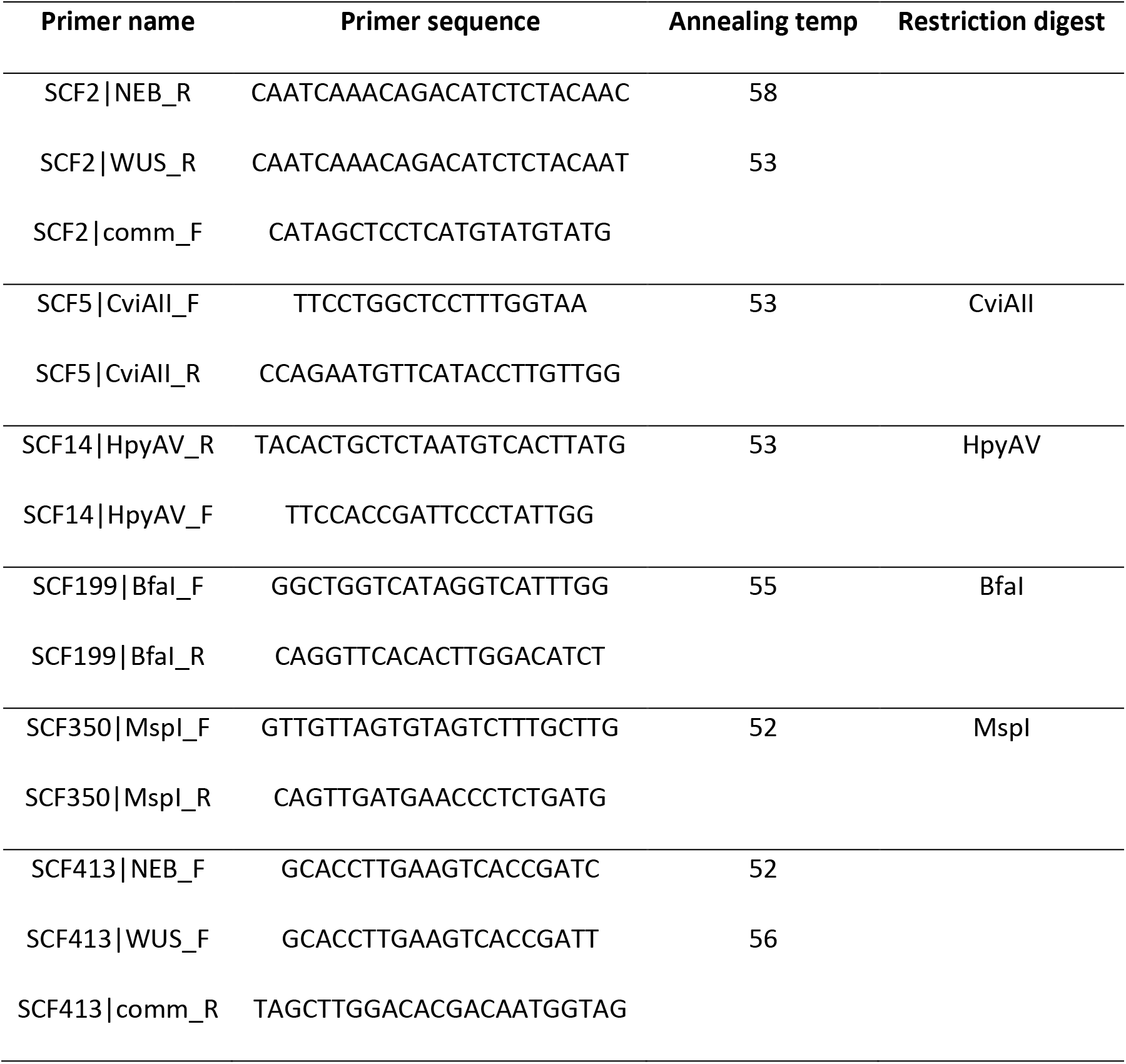
Parent-specific primers within putative quantitative trait loci for tembotrione resistance, their associated annealing temperatures, and restriction enzymes.

## 3 RESULTS

Tembotrione treatment of vegetative clones from the parental lines was used to select sensitive and resistant plants to generate the pseudo-F_2_ mapping population. Characteristic tembotrione bleaching was observed throughout the new leaves of selected WUS plants in response to 0.92 g ai ha^-1^, while no injury phenotypes were observed in selected NEB plants, even though they were screened with a 15-times higher rate (data not shown). The F_1_ progeny derived from the RxS cross were challenged with a rate titration of tembotrione (Figure 1). No segregation (data not shown) and a dominance inheritance pattern within the F_1_ was observed. An R/S ratio of 15.0 (RxR) and 21.1 (RxS) was observed (Table 2), though the model generated for the RxR population was not significantly different from that of the RxS population (p = 0.20). Based on these dose-response curves, a delimiting rate of 11.5 g tembotrione ha^-1^ was selected for screening the pseudo-F_2_ population. This screening rate is notably lower than the standard field use rate of approximately 100 g ai ha=^1^, which is not uncommon for greenhouse dose-response trials.

**Figure 1.**
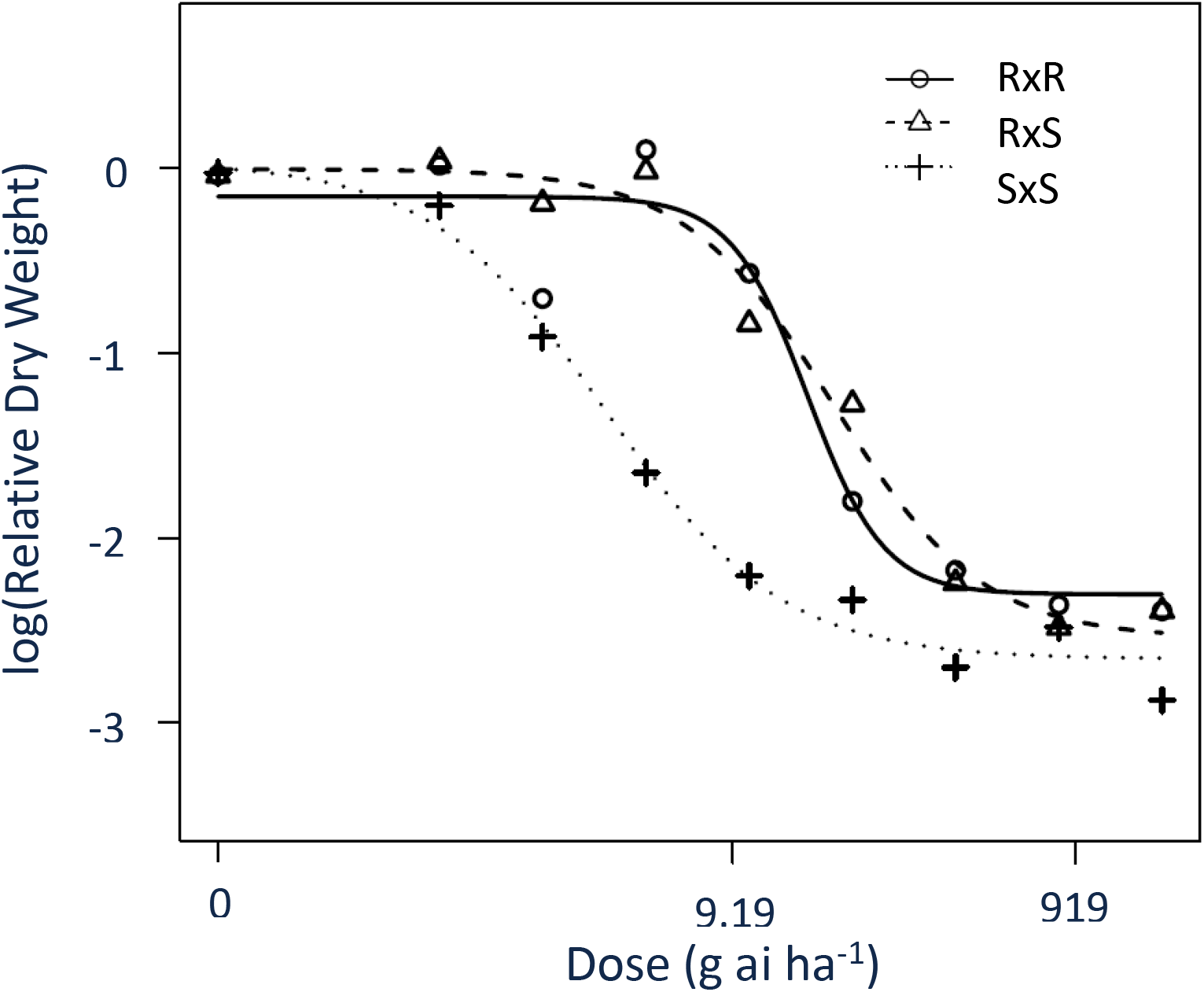
Dose response models of RxR, RxS, and SxS F_1_ populations derived from NEB (R) and WUS (S). Relative biomass was calculated against the average response to 0 g ai ha^-1^ tembotrione 21 days after application.

**Table 2.**
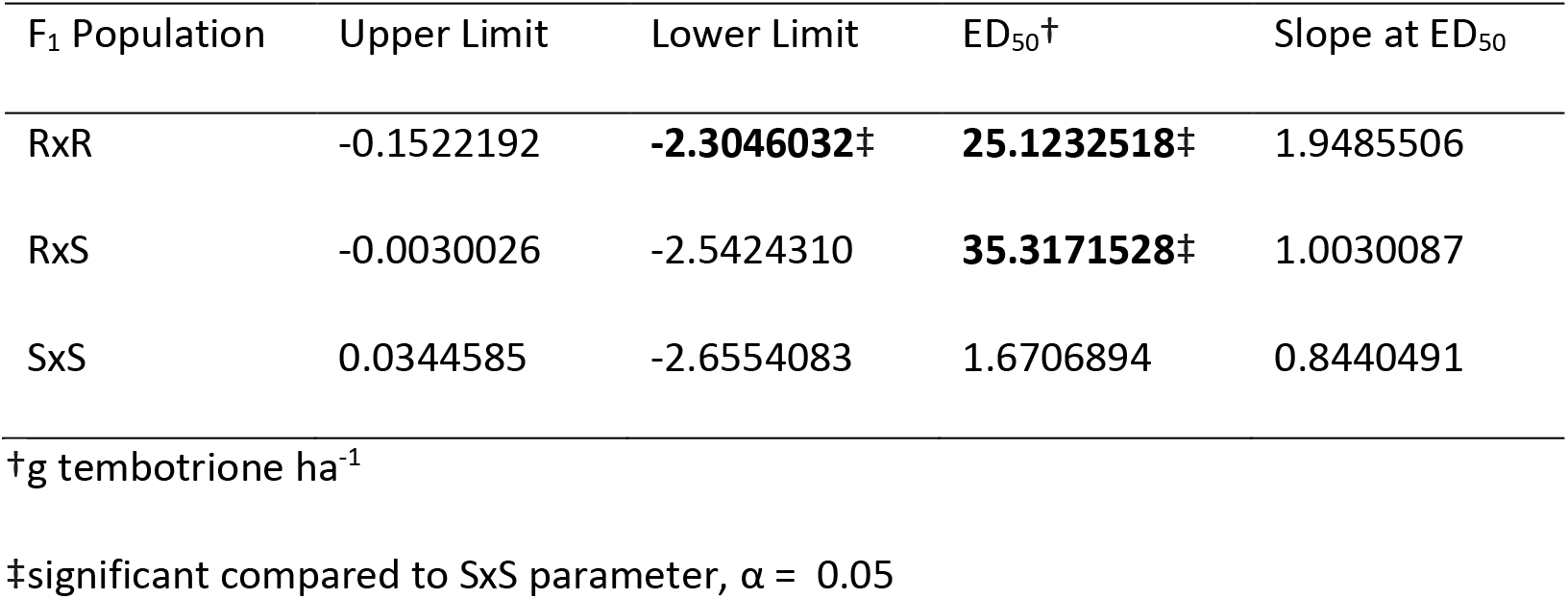
Parameter estimates for dose-response analysis of resistant (RxR), F_1_ (RxS), and sensitive (SxS) populations in response to tembotrione application, based on above ground dry weight 21 days after application.

The two screening runs of the pseudo-F_2_ population could not be combined statistically and are presented separately in Figure 2. For both runs, a series of two peaks were observed, with one each in proximity of the mean phenotypic values of the SxS and RxR populations. Broad-sense heritability was calculated to be 0.556, suggesting a moderately heritable trait.

**Figure 2.**
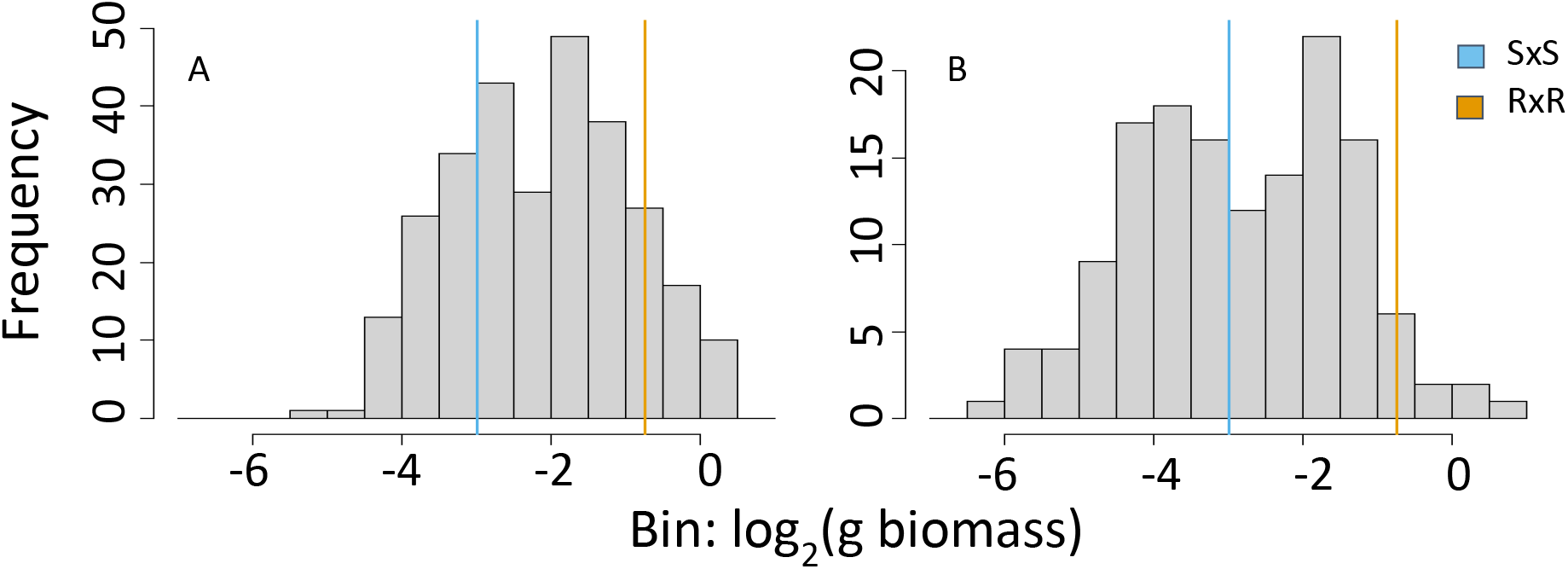
Distribution of dry weights of the pseudo-F_2_ population in response to tembotrione, 21 days after application. Mean phenotypes of parental populations (RxR, SxS) are shown in colored lines. Replication A and B were performed in 2019 and 2020, respectively.

Bulk segregant analysis was conducted on computationally binned pseudo-F_2_ individuals. Five putative QTL were identified through this analysis, each on different chromosomes (Table 3). Except for the Scaffold 5 QTL, each QTL spanned a single contig of the genomic assembly. The putative QTLs ranged in size from 11 Kbp to 522 Kbp. Molecular markers were developed within each contig of the putative QTLs and screened for association with increased biomass 21 days after herbicide application. At the tested sample size (N = 276), only two putative QTL could be confirmed, one on scaffold 4 and one on scaffold 12. The QTL on scaffold 4 appeared dominant in nature (Supplementary Figure 1), and the RR and RS bins were combined (Figure 3A), with an effect size of 0.09 g biomass 21 days after application. The QTL on scaffold 12 appears additive (Figure 3B), with an effect size of 0.06 g (per allele). In Figure 3C, the combined effect of both QTLs is shown. The absence of resistance alleles at both QTLs resulted in a bin of plants statistically similar to the sensitive control population in tembotrione response. The presence of more resistance alleles resulted in a stepwise increase in the mean phenotype of the given bin, and the R_+RR (Scaffold 4 + Scaffold 12) genotype was statistically similar to the RxR population. Interestingly, the R_+RS genotypic bin, largely corresponding to the genotype of the F_1_ population, was statistically different from both the SxS and RxR controls. In contrast, the F_1_ inheritance study indicated a dominant trait. Representative plants of the pseudo-F_2_ population, run A, are shown in Supplemental Figure 2.

**Table 3.**
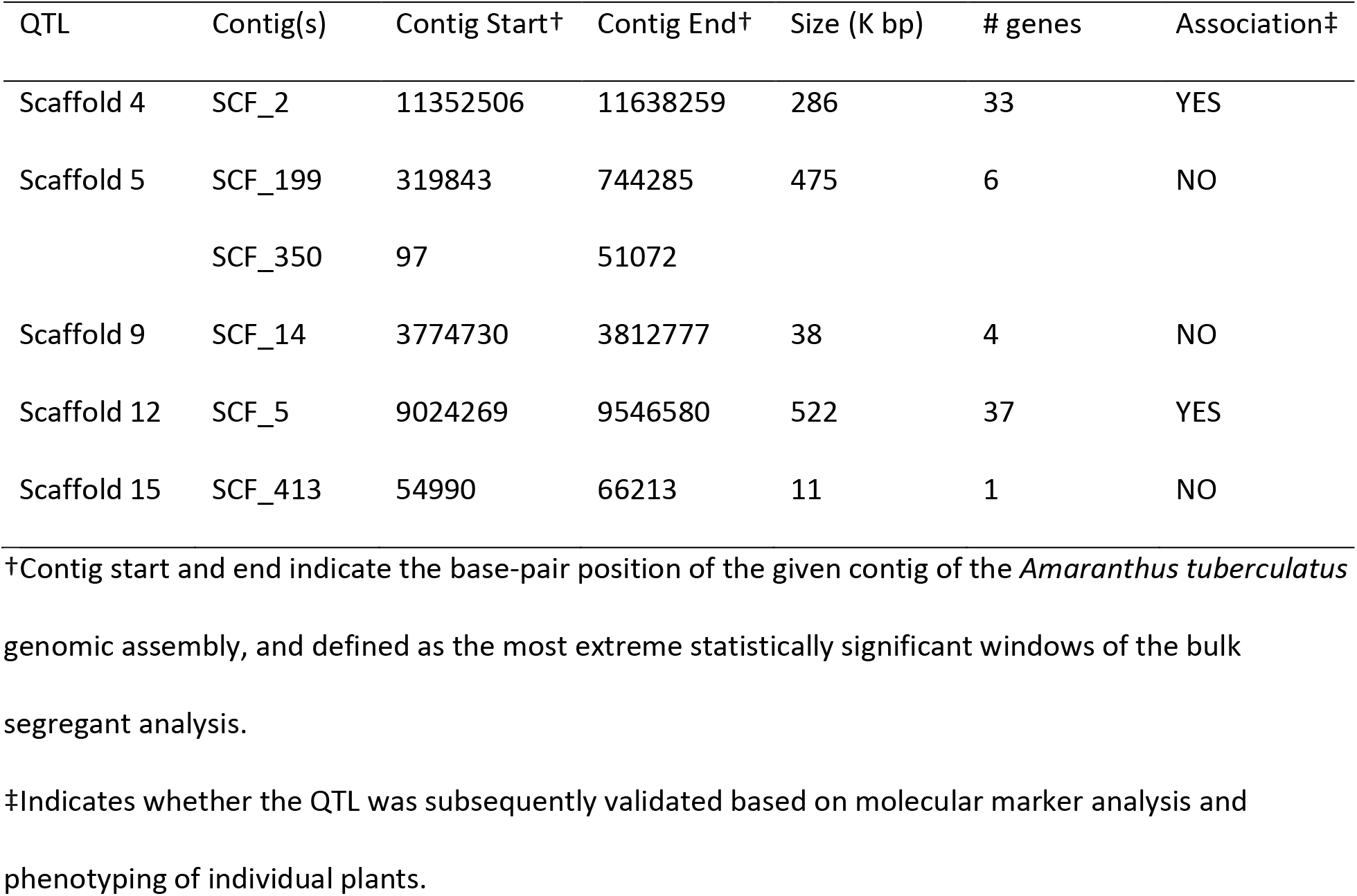
Putative quantitative trait loci (QTL) identified through bulk segregant analysis and association with tembotrione resistance in additional pseudo-F_2_ plants.

| QTL | Contig(s) | Contig Start<sup>†</sup> | Contig End<sup>†</sup> | Size (K bp) | # genes | Association<sup>‡</sup> |
| --- | --- | --- | --- | --- | --- | --- |
| Scaffold 4 | SCF_2 | 11352506 | 11638259 | 286 | 33 | YES |
| Scaffold 5 | SCF_199 | 319843 | 744285 | 475 | 6 | NO |
|  | SCF_350 | 97 | 51072 |  |  |  |
| Scaffold 9 | SCF_14 | 3774730 | 3812777 | 38 | 4 | NO |
| Scaffold 12 | SCF_5 | 9024269 | 9546580 | 522 | 37 | YES |
| Scaffold 15 | SCF_413 | 54990 | 66213 | 11 | 1 | NO |

<sup>†</sup>Contig start and end indicate the base-pair position of the given contig of the *Amaranthus tuberculatus* genomic assembly, and defined as the most extreme statistically significant windows of the bulk segregant analysis.
<sup>‡</sup>Indicates whether the QTL was subsequently validated based on molecular marker analysis and phenotyping of individual plants.

**Figure 3.**
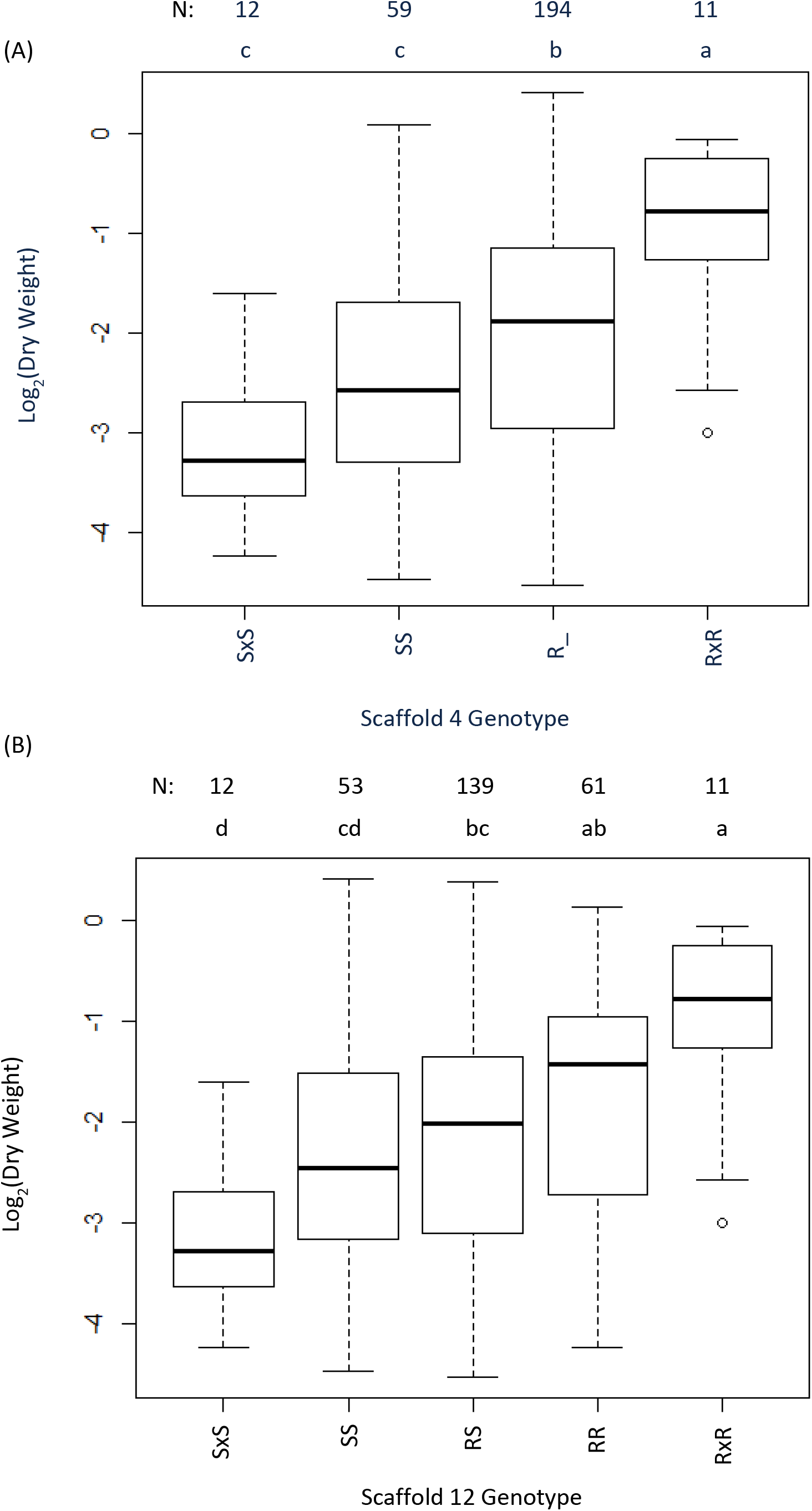
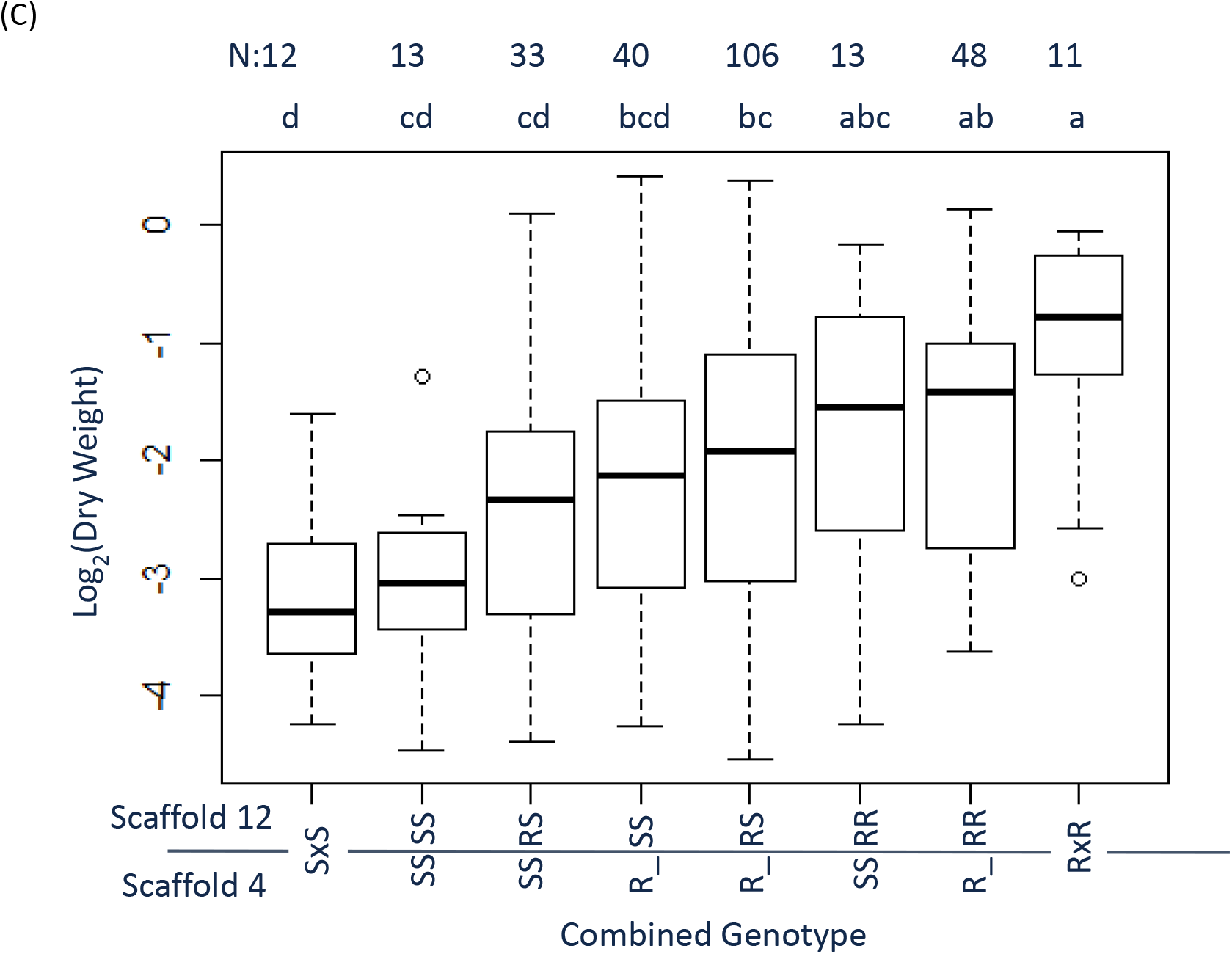
Phenotypic responses of tembotrione-sensitive (SxS) and resistant (RxR) parents and of pseudo-F_2_ individuals binned by genotype (parental allele inheritance at quantitative trait loci (QTL) within Scaffold 4 and 12). (A) Samples binned by Scaffold 4 genotype, which follows a dominant inheritance pattern. (B) Samples binned by Scaffold 12 genotype, which follows an additive inheritance pattern. (C) Samples were binned by both Scaffold 4 and Scaffold 12 genotype. Because the Scaffold 12 QTL appeared to be dominant, plants heterozygous and homozygous for the R allele were combined (R_) in panels A and C.

A total of 70 gene models were observed within the boundaries of the two validated QTL, 33 on Scaffold 4 and 37 on Scaffold 12. Thirty-three gene models were annotated, however no cytochrome P450s genes, the canonical gene for HPPD-inhibitor resistance, were observed. Annotations associated with cell wall traits, metabolism, and transcription factors were observed (Supplementary File 1).

## 4 DISCUSSION

The continued evolution of weeds under the selection pressures of agricultural management has profound implications for food security^32^. The identification and characterization of heritable factors that result in herbicide resistance provide vital knowledge into how plants adapt to abiotic stimuli. The adoption of next generation sequencing technologies and the development of impactful genomic resources for problematic weed species is critical to deciphering the underlying mechanisms that drive the response to selection^33–35^. Herbicide resistance is frequently delimited into target-site and non-target-site mechanisms, based on the involvement of proteins with which the herbicide interacts to elicit plant death. Because the target-site gene(s) are frequently known and conserved within and across plant species, the identification of the specific heritable factors is well reported across literature^36,37^. In contrast, non-target-site mechanisms are largely approached from a physiological perspective, where higher-level analyses such as metabolic profiling and differential gene expression are conducted^38^. While these approaches have identified numerous candidate genes, and led to the functional validation of others, these approaches do not capture the underlying genetic architecture of the queried herbicide-resistance traits. Here, within the weed species *A. tuberculatus*, we present the first QTL-mapping study to characterize the genetic architecture of herbicide resistance.

Efforts to investigate the evolutionary origins of herbicide resistance have utilized population genomics screens in unstructured populations^19,39,40^. In the case of *Ipomoea purpurea* L. Roth, five genomic regions under selection to glyphosate were identified through the comparison of 80 samples from eight populations, while Kreiner et al investigated 162 samples across 21 collection sites and several natural areas^19^. In each case, the genetic architecture underlying the resistance traits was complex, and numerous candidate genes were proposed. Similarly, we observed a complex genetic architecture controlling HPPD-inhibitor resistance in *A. tuberculatus* through our QTL-mapping approach. Both GWAS and QTL mapping approaches provide critical insights into the evolution of herbicide resistance in weedy species. Both approaches are complementary and should be used in concert^41^. While association mapping techniques are often favored due to high resolution and the ease of obtaining non-structured populations, the structured populations of a linkage mapping study provide ideal material for subsequent physiological and functional analysis^42^.

Previously, an RNA-seq experiment was conducted on pseudo-F_2_ segregants of the NEB population revealing 115 differentially expressed transcripts and 268 differentially expressed genes^23^. Several of these were annotated as metabolism genes, including a glucosyl-transferase and an ABC transporter. Interestingly, no differentially expressed genes or transcripts overlapped with the QTL identified in this study. The transcriptome project was derived from different parent plants, therefore a different pseudo-F_2_ population, and, as such, the presence of multiple resistance mechanisms within the NEB population would confound this comparison. In the absence of functional validation, QTL are specific to the population in which they were identified. Ideally, the two studies would have been conducted on the same pseudo-F_2_ population. In addition, these two studies investigated the HPPD-inhibitor phenotype at different regulatory levels. Gene expression, as captured through transcriptome profiling, is modulated through both proximal and distal elements. Furthermore, modification of transcription factors allows for trans-acting modulation of gene expression. Therefore, we suggest that the differential regulation observed by Giacomini et al is the result of regulation by distal factors^23^. In a different population, Kohlhase et al identified Scaffold 15 as enriched for SNP associated with HPPD-inhibitor resistance in *A. tuberculatus^15^*. The QTL found in this study were not observed on scaffold 15, suggesting resistance to HPPD-inhibitors has evolved independently within *A. tuberculatus*.

Heritability and gene action are important considerations for the spread of herbicide resistance. In this study, the HPPD-inhibitor resistance was moderately heritable, though resulted in a 15-21 fold rate shift to achieve 50% control. Indeed, while heritability is a key factor in the selection response, labeled application rates in a field setting do not vary on the scale of orders of magnitude. A trait with low heritability, though with a sufficient effect size, could result in resistance evolution over a single cycle of selection. In the case of HPPD-inhibitor resistance in the NEB population, moderate heritability may not be unexpected. Phenotypic variation is clearly observable within progeny from RxR and SxS crosses, indicating the presence of environmental variation. Furthermore, the observed segregation suggests that maternal effects are minimal, if present. Additionally, while homozygous for the two major QTL identified in the study, small effect variants segregating within the source NEB population, which could not be captured at the tested sample size, could also contribute to the observed variation. Due to the lack of doubled-haploid technology, the dioecious nature of *A. tuberculatus*, and severe inbreeding depression, the development of inbred material was impractical.

Effect sizes of Scaffold 4 and 12 QTL were calculated to be 0.09 g biomass and 0.06 g biomass per allele, respectively, 21 days after application. Conventionally, effect size is expressed through an R/S ratio determined through dose response analysis, since the value can be heuristically associated with in-field resistance. While dose-response modelling is widely used to account for environmental variation^43^, delimiting rate screening remains the mainstay of segregation studies. Here, we present our effect sizes as the biomass shifts observed between genotypic bins in the pseudo-F_2_ generation, as a delimiting rate screening was used. The stabilization of each QTL individually into lines, such as near-isogenic lines, would enable accurate prediction of effect sizes as an R/S ratio, which possesses real-world applications. In contrast to the dominant gene action suggested by the F_1_ inheritance, an additive response was observed for the Scaffold 12 QTL. Furthermore, the R_+RS genotypic bin was significantly differentiated from either parental population, suggesting a partially additive trait inheritance (Figure 3C). As the F_1_ and pseudo-F_2_ trials were separated by time, environmental variation between the experiments could account for the difference in observed inheritance. In addition, the additive nature of the Scaffold 12 QTL could have been masked through the genetic background of the NEB population. Conventionally, gene action for herbicide resistance is based on the response of F_1_ populations. Alternatively, linkage mapping in the pseudo-F_2_ generation investigates gene action after independent assortment between the sensitive and resistant backgrounds, which is preferable. From these observations, we propose that linkage mapping and other associated techniques are ideal to determine gene action. Alternatively, the development of resistant and sensitive lines from a structured population should be used to ensure a common genetic background.

Reports and subsequent characterization of HPPD-inhibitor resistance has largely focused on metabolic profiling. Indeed, numerous studies have pointed towards cytochrome P450s as being pivotal for resistance to the herbicide group^8–15^. Recently, Jacobs et al reported a tight association between resistance to atrazine and a component of a complex HPPD-inhibitor resistance phenotype in *A. tuberculatus^44^*. While atrazine resistance in the tested population is mediated by a glutathione *S*-transferase (Ma et al 2013), the mechanism of resistance to HPPD-inhibitors is unknown. Intriguingly, no cytochrome P450s were observed within the regions delimited by either QTL. Linkage mapping approaches identify regions of the genome associated with the causal variant, but this variant does not necessarily have to be a gene whose product is directly associated with the physiological basis of resistance. A distal regulatory element, either *cis*- or *trans*-acting, could be the causal heritable factor. Alternatively, resistance could be due to factors other than cytochrome P450s, such as reduced translocation. Physiological evaluation of the resistance phenotypes will prove critical in understanding the functional relationship between the identified QTL and the herbicide-resistant phenotypes.

In the future, we plan on conducting metabolic profiling to gain insight into the physiological basis of the HPPD-inhibitor resistant trait in the NEB population. Metabolic profiling, and other approaches to investigate the physiological basis of resistance, are critical to the identification of causal genes that underlay the resistance phenotype. Genetic and physiological approaches are naturally synergistic to achieving this goal. Genetic analysis, including linkage mapping, transcriptome profiling, and association mapping, often produce lists of genes distributed amongst many protein families. Conversely, physiological approaches frequently identify key gene families which underlie the trait of interest. A combined approach can rapidly generate a small subset of candidate genes for functional validation. In this study, we have begun to unravel a multi-gene herbicide resistance trait. Ideally, physiological analysis would be conducted on near-isogenic lines to minimize the background effects present between different populations to focus on the specific QTL of interest. Therefore, stabilizing each QTL into lines will both allow rigorous dose-response analysis and metabolic profiling to be conducted.

## 5 CONCLUSION

We report the first QTL mapping study for herbicide resistance in weedy species. HPPD-inhibitor resistance in the NEB population is a complex and largely dominant trait. Two large-effect QTL, on Scaffold 4 and Scaffold 12, for HPPD-inhibitor resistance were identified and validated within this population.

## ACKNOWLEDGEMENTS

This research was supported by a grant from Bayer AG, Division of Crop Science. No other conflict of interest is declared.

