## Supplemental Figure 1 for "Genetic architecture underlying HPPD-inhibitor resistance in a Nebraska *Amaranthus tuberculatus* population"

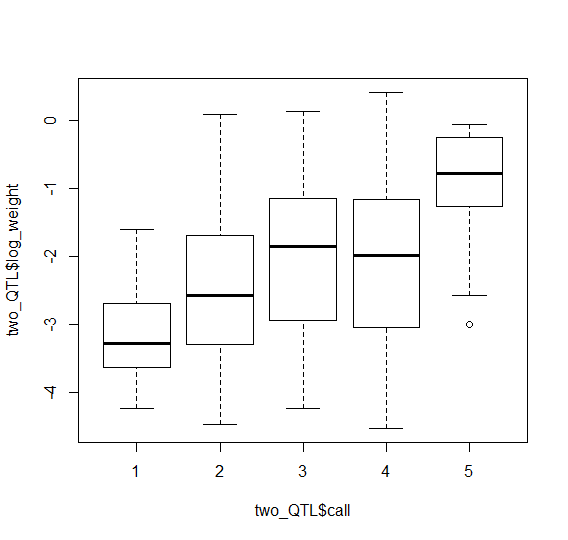

c	bc	b	ab	a

SxS

RxR

RR

RS

SS

Log_2_(Dry Weight (g))

Scaffold 4 Genotype

N: 12	59	142	52	11

Supplementary Figure 1. Phenotypic response of tembotrione-sensitive (SxS) and resistant (RxR) parents and of pseudo-F_2_ individuals 21 days after application binned by genotype.
