## Supplemental Figure 2 for "Genetic architecture underlying HPPD-inhibitor resistance in a Nebraska *Amaranthus tuberculatus* population"

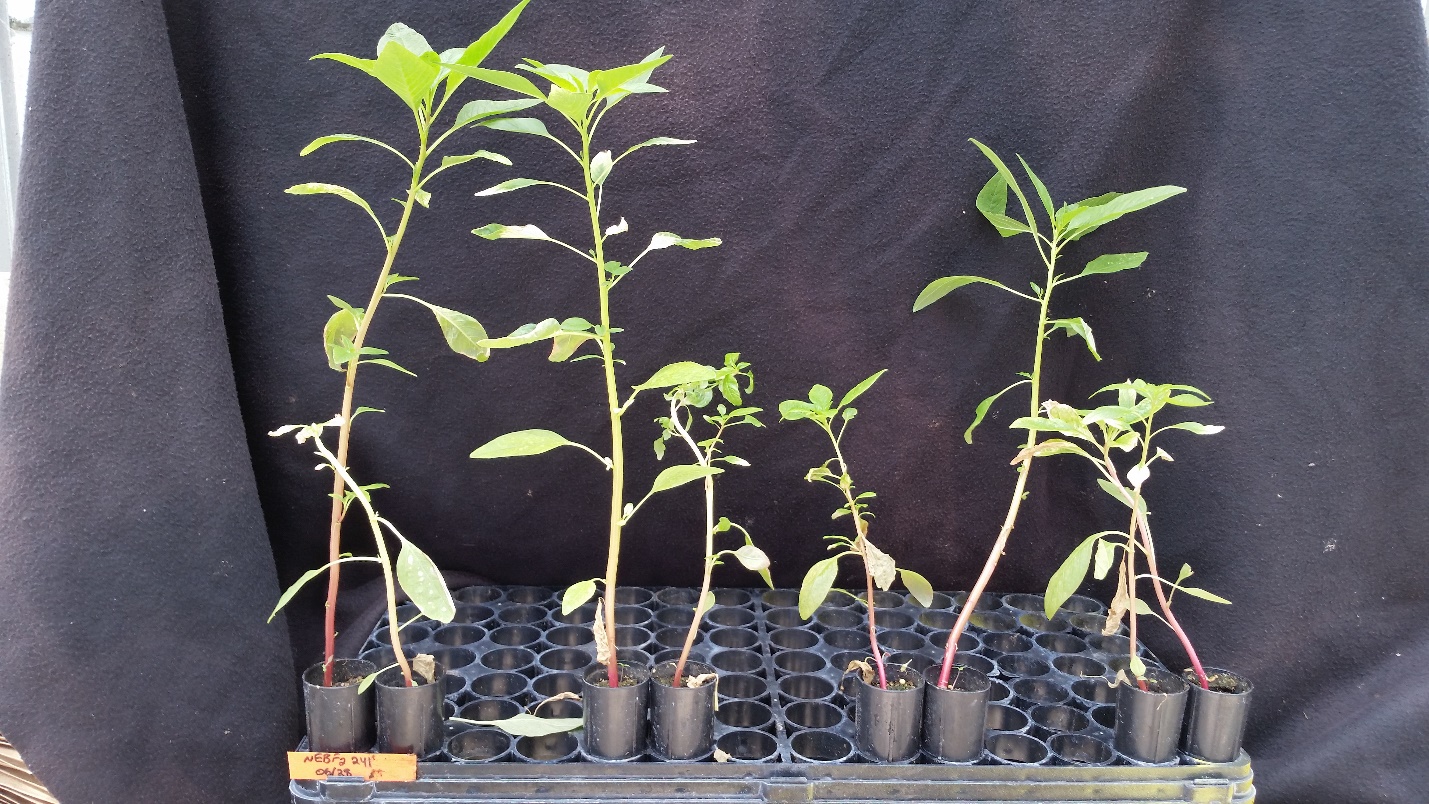

Supplementary Figure 2. Representative phenotypic response of pseudo-F_2_ individuals 21 days after application with tembotrione. Genotypes from left to right (Scaffold 4/Scaffold 12): RS/RS, SS/RS, RR/RR, RR/SS.
